# Re-awakening the brain: Forcing transitions in disorders of consciousness by external *in silico* perturbation

**DOI:** 10.1101/2023.07.17.549269

**Authors:** Paulina Clara Dagnino, Anira Escrichs, Ane López-González, Olivia Gosseries, Jitka Annen, Yonatan Sanz Perl, Morten L. Kringelbach, Steven Laureys, Gustavo Deco

**Author notes:** Correspondig authors (P.D), (A.E), (G.D). These authors contributed equally to this work (Paulina Clara Dagnino, Anira Escrichs). These authors share senior authorship (Steven Laureys, Gustavo Deco).

## Abstract

A fundamental challenge in neuroscience is accurately defining brain states and predicting how and where to perturb the brain to force a transition. The ability to promote a transition from one brain state to another by externally driven stimulation could significantly impact rehabilitation and treatments for patients suffering from complex brain injury cases. Thus, it is crucial to find therapeutic interventions able to re-balance the dynamics of brain disorders towards more healthy regimes. Here, we investigated resting-state fMRI data of patients suffering from disorders of consciousness (DoC) after coma (minimally conscious and unresponsive wakefulness states) and healthy controls. We applied model-free and model-based approaches to help elucidate the underlying brain mechanisms of patients with DoC. The model-free approach allowed us to characterize brain states in DoC and healthy controls as a probabilistic metastable substate (PMS) space. The PMS of each group was characterized by a repertoire of unique patterns (i.e., metastable substates) with different probabilities of occurrence. In the model-based approach, we adjusted the PMS of each DoC group to a causal whole-brain model. This allowed us to explore optimal strategies for promoting a transition to the PMS of the control group by applying off-line *in silico* probing. Furthermore, this approach enabled us to evaluate the impact of all possible local perturbations in terms of their global effects and sensitivity to stimulation, which is a biomarker providing a deeper understanding of the mechanisms underlying DoC. Our results show that transitions from DoC to more healthy regimes were obtained in a synchronous protocol, in which areas from the motor and subcortical networks were the most sensitive to perturbation. This motivates further work to continue understanding brain function and treatments of disorders of consciousness by external stimulation.

**Author summary:** We studied disorders of consciousness by defining a brain state as a repertoire of metastable substates with different probabilities of occurrence. We created whole-brain computational models of DoC to uncover the causal mechanisms underlying recovery. These models allowed us to transition from DoC to a control healthy state by studying the effects of artificial individual local perturbations under different protocol regimes. We demonstrated successful transitions in the synchronization protocol and showed that the most sensitive areas were located in the motor network and subcortical regions. We believe this could be very valuable for developing clinical treatments and has a great deal for future therapies.

## Introduction

The brain is a dynamical, complex, and self-organized system with spontaneous activity emerging from non-linear interactions of billions of neurons (Sporns, 2011). This gives rise to an ample discrete repertoire of metastable patterns (i.e., substates) around critical points between order and chaos (Cabral et al., 2017b; Deco et al., 2017c). The lifetimes and stabilities of specific substates govern the dynamics of a particular brain state (Deco and Kringelbach, 2016; Tognoli and Kelso, 2014). Current research is increasing our understanding of the causal dynamics underlying many different brain states, such as wakefulness, sleep, anesthesia, and disorders of consciousness (DoC). Nevertheless, such mechanisms still remain elusive and a deeper comprehension would facilitate the design of novel interventions for brain disorders and possibly for the loss of consciousness like coma. Recently, directly perturbing the brain *in silico* has been proposed and investigated as a possible intervention that could contribute to a deep understanding of the dynamical mechanisms of brain states in health and disease (Escrichs et al., 2022; Vohryzek et al., 2022a; Deco et al., 2019). Furthermore, such perturbations could be used to force transitions between different brain states in a translational clinical context, for example, to promote transitions from brain disorders to health (Fox et al., 2014; Thibaut et al., 2014; Schiff et al., 2007).

A healthy brain relies on the brain’s flexibility and capacity to integrate information and maintain rich dynamics in an evolving environment across time and space (Deco et al., 2015). By contrast, brain disorders present disruptions in the normal range of brain activity (Du et al., 2018). In the specific clinical domain of DoC, it has been found that their characteristic brain patterns present disruptions of long-range cortical correlations typical in a healthy state (Demertzi et al., 2019). Such post-coma states are distinguished into the minimally conscious state (MCS) and unresponsive wakefulness syndrome (UWS). The former MCS is identified when patients are awake and respond with limited awareness, and the latter UWS corresponds to patients who do not respond to stimulation in a conscious manner (Giacino et al., 2018, 2002). DoC patients present lower flexibility and efficiency of information processing and a limited broadcast of information, which coexists with a reduced neural propagation and responsiveness to events (Panda et al., 2023). Furthermore, UWS shows reduced metastability and repertoire of functional networks in comparison to MCS (Panda et al., 2022).

In recent years, different definitions of brain states have been proposed using empirical neuroimaging and electrophysiological data. Approaches based on functional magnetic resonance imaging (fMRI) have implemented static analysis such as long-range temporal dependence via Hurst exponent (Tagliazzucchi et al., 2013) and attractors between brain regions (Gu et al., 2018; Deco and Jirsa, 2012). Considering brain activity is multi-dimensional and ever-changing, these approaches have been further examined from a more realistic and richer viewpoint considering brain dynamics, which reveals the different brain patterns evolving during a scanning period (Escrichs et al., 2021b; Sanz Perl et al., 2022; Deco and Kringelbach, 2020; Preti et al., 2017; Hansen et al., 2014; Allen et al., 2014; Hutchison et al., 2013). Nevertheless, a universal, formal, robust, and quantitative definition of brain states, and a deep comprehension of the effects of perturbations to force recovery, remains unknown (Deco et al., 2019, 2017c). Stemming from recent progress in these areas, and given the difficulty of predicting the final collective emergent activity even if the building blocks are known (Deco et al., 2019), we could still benefit from a better understanding of brain dynamics and optimal strategies for a recovery towards healthy brain states (Vohryzek et al., 2022a; Escrichs et al., 2022; Kringelbach and Deco, 2020; Edlow et al., 2020; Deco et al., 2017a; Keilholz et al., 2017).

There is a long tradition of perturbative approaches for brain research. Clinical techniques for stimulation exist, such as the non-invasive transcranial direct current stimulation (tDCS) (Knotkova et al., 2019; Ruffini et al., 2018; Siebner et al., 2009) and transcranial magnetic stimulation (TMS) (Litvak et al., 2007; Pascual-Leone, 1999), and the minimally invasive technique deep brain stimulation (DBS) (Mohseni et al., 2012; Kringelbach et al., 2007). Still, research in lesioned humans is rare, only undertaken when the disease is severe, and accompanied by ethical constraints (Deco and Kringelbach, 2017; Clausen, 2010). Massimini and colleagues developed the perturbational complexity index (PCI), which has been used to distinguish brain states by calculating the lempel-ziv complexity from the electroencephalography response to TMS perturbation (Casarotto et al., 2016; Casali et al., 2013; Massimini et al., 2009). The PCI measures the perturbation-elicited variations in intrinsic global brain activity and has shown to be successful in distinguishing between awake vs. sleep, awake vs. anesthesia, and MCS vs. UWS (Casarotto et al., 2016; Casali et al., 2013; Ferrarelli et al., 2010; Massimini et al., 2009). However, given the ethical restrictions of empirical neurostimulation approaches, causal whole-brain models based on *in silico* perturbation protocols are fundamental to understanding the underlying mechanisms of brain dynamics (Escrichs et al., 2022). This promising tool allows experimenting in unprecedented unlimited scenarios (e.g., perturbing one brain area at a time) without exposing real brains (Kringelbach and Deco, 2020; Breakspear, 2017; Deco et al., 2015; Deco and Kringelbach, 2014).

Recently, Deco et al. (2019) proposed the awakening framework that consists of model-free and model-based approaches to force transitions from deep sleep to awake. In particular, the model-free approach based on Leading Eigenvector Dynamics Analysis (LEiDA) (Cabral et al., 2017b) uses the concept of metastability, defined as the characteristic of a system to maintain an equilibrium in a temporal window although being slightly perturbed (Freyer et al., 2012; Kelso, 2012; Freyer et al., 2011). The nature, duration, and arrangement of existent metastable substates (i.e., patterns) give rise to a probabilistic metastable substate (PMS) space typifying each brain state (Deco et al., 2017c). LEiDA has been shown to be robust and successful in identifying brain states in healthy aging (Escrichs et al., 2021a; Cabral et al., 2017b), depression (Figueroa et al., 2019) and different states of consciousness (Kringelbach and Deco, 2020; Lord et al., 2019; Deco et al., 2019). The model-based approach consists in building whole-brain models composed of a network of coupled local nodes (Botvinik-Nezer et al., 2020; Deco et al., 2019) to simulate the empirical PMS and perturb the resulting PMS model to force the transition to a desired control state. This elegant framework has been extended to promote transitions from aging (Escrichs et al., 2022), patients with depression (Vohryzek et al., 2022b) and schizophrenia (Mana et al., 2023) towards more healthy regimes.

Here, we aimed to study the dynamical complexity and causal mechanisms of brain activity in DoC by using the aforementioned framework. Firstly, we applied LEiDA to define the PMS of DoC patients and healthy controls. Secondly, we built Hopf whole-brain models fitted and optimized to the empirical PMS of DoC at the bifurcation point, representing a state of criticality in which the two regimes (oscillatory and noisy) cannot be differentiated. This generative whole-brain model linked structural anatomy with functional dynamics on the basis of effective connectivity (Deco et al., 2019). Finally, we applied off-line *in silico* external unilateral and localized probing to force the transition from the PMS obtained in MCS and UWS, separately, to the PMS of healthy controls. In this way, employing offline *in silico* probing, we could evaluate candidate regions for stimulation aiming to recover DoC patients. Nevertheless, this innovative approach not only allowed us to assess the effects of all potential local perturbations but also provided valuable insights into their mechanistic global effects and sensitivity to stimulation.

## Materials and Methods

### Ethics statement

The study was approved by the Ethics Committee of the Faculty of Medicine of the University of Liège according to the Helsinki Declaration on ethical research. Written informed consent was obtained from controls and the patients’ legal surrogates.

### Participants

A total of 23 controls and 46 non-sedated patients with DoC were selected from a dataset previously described in Escrichs et al. (2021b); López-González et al. (2021); Demertzi et al. (2019). Trained clinicians carried out the clinical assessment and Coma Recovery Scale-Revised (CRS-R) scoring to determine the patients’ state of consciousness. The CRS-R diagnosis was made after at least 5 CRS-R, and the highest level of consciousness was taken as the final diagnosis, which was also confirmed using positron emission tomography (PET) (i.e., patients in MCS presented a relatively preserved metabolism in the frontoparietal network, whilst patients with UWS had a bilateral hypometabolism in this network). Thus, 30 patients in MCS and 16 in UWS were included.

### MRI Data Acquisition

MRI data were acquired on a 3T Siemens TIM Trio scanner (Siemens Inc, Munich, Germany). Resting-state fMRI data were obtained using a gradient echo-planar imaging (EPI) sequence (300 volumes, 32 transversal slices, TR= 2000 ms, TE=30 ms, flip angle = 78°, voxel size = 3×3×3 mm, FOV = 192 mm). After fMRI acquisition, a structural T1 magnetization-prepared rapid gradient-echo (MPRAGE) sequence was acquired (120 slices, TR = 2300 ms, voxel size = 1.0×1.0×1.2 mm, flip angle = 9°, FOV = 256 mm). Finally, diffusion-weighted MRI (DWI) was acquired with 64 directions (b-value =1,000 s/mm^2^, voxel size = 1.8 × 1.8 × 3.3 mm^3^, FOV = 230 × 230 mm^2^, TR/TE= 5,700/87 ms, 45 transverse slices, 128 × 128 voxel matrix) preceded by a single b0 image.

### Resting state fMRI preprocessing

The pre-processing of resting-state fMRI data was performed using MELODIC (Multivariate Exploratory Linear Optimized Decomposition into Independent Components) version 3.14 (Beckmann and Smith, 2004) from FMRIB’s Software Library (FSL, http://fsl.fmrib.ox.ac.uk/fsl) as described in our previous studies (Escrichs et al., 2021b; López-González et al., 2021). The following steps were performed: discarding the first 5 volumes, motion correction motion using MCFLIRT (Jenkinson et al., 2002), non-brain removal using BET (Brain Extraction Tool) (Smith, 2002), spatial smoothing with a 5 mm Gaussian Kernel, rigid-body registration, high pass filter (with a cutoff of 100 s) and single-session Independent Component Analysis (ICA) with automatic dimensionality estimation. Then, noise components and lesions-driven artifacts (for patients) were manually classified and removed for each subject by looking at the spatial map, time series, and power spectrum (Griffanti et al., 2017; Salimi-Khorshidi et al., 2014) using FIX (FMRIB’s ICA-based X-noiseifier) (Griffanti et al., 2014). Finally, FSL tools were used to co-register the images and extract the time series between 214 cortical and subcortical brain areas for each subject in MNI space from the Shen resting-state atlas (without the cerebellum) (Shen et al., 2013).

### Probabilistic Tractography preprocessing

A whole-brain structural connectivity (SC) matrix was computed for each subject of the control group and then averaged in a two-step process as described in previous studies (Muthuraman et al., 2016; Cao et al., 2013; Gong et al., 2009). We used the resting-state atlas mentioned above to create a structural connectome in each individual’s diffusion native space. In brief, DICOM images were converted to Neuroimaging Informatics Technology Initiative (NIfTI) using dcm2nii (www.nitrc.org/projects/dcm2nii). The b0 image in DTI native space was co-registered to the T1 structural image by using FLIRT (Jenkinson and Smith, 20001). Then, the T1 structural image was co-registered to the standard space by using FLIRT and FNIRT (Andersson et al., 2007; Jenkinson and Smith, 20001). The transformations were inverted and applied to warp the resting-state atlas from MNI space to the native diffusion space by applying a nearest-neighbor interpolation method. Analysis of diffusion images was performed using the processing pipeline of the FMRIB’s Diffusion Toolbox (FDT) in FMRIB’s Software Library www.fmrib.ox.ac.uk/fsl. Non-brain tissues were extracted using Brain Extraction Tool (BET) (Smith, 2002), eddy current-induced distortions and head movements were corrected using eddy correct tool (Andersson and Sotiropoulos, 2016), and the gradient matrix was reoriented to correct for subject motion (Leemans and Jones, 2009). Then, Crossing Fibres were modeled using the default BEDPOSTX parameters, and the probability of multi-fibre orientations was computed to improve the sensitivity of non-dominant fibre populations (Behrens et al., 2007, 2003). Probabilistic Tractography was performed in native diffusion space using the default parameters of PROBTRACKX (Behrens et al., 2007, 2003). The connectivity probability to each of the other 214 brain areas was estimated for each brain area as the total proportion of sampled fibres in all voxels in the brain area *n* that reached any voxel in the brain area *p*. Given that Human Difussion Tensor Imaging (DTI) does not capture directionality, the *SC*_*np*_ matrix was symmetrized by computing its transpose *SC*_*pn*_ and averaging both matrices. Finally, to obtain the structural probability matrix, the value of each brain area was divided by its corresponding number of generated tracts.

### Leading Eigenvector Dynamics Analysis (LEiDA)

This first step aims to define the empirical brain states from a quantitative point of view, defined as a conjunction of substates, applying LEiDA method (Cabral et al., 2017b) as schematized in **Figure 1a**. For all subjects in all states, the blood oxygenation level-dependent (BOLD) time series of each brain area of the parcellation were filtered in the range 0.04-0.07 Hz and Hilbert-transformed to obtain the evolution of phase of the time series. A BOLD phase coherence matrix dFC(*t*) was then calculated at any given repetition time (TR) between each brain area pair *n* and *p* by calculating the cosine of the phase difference as:

**Figure 1:**
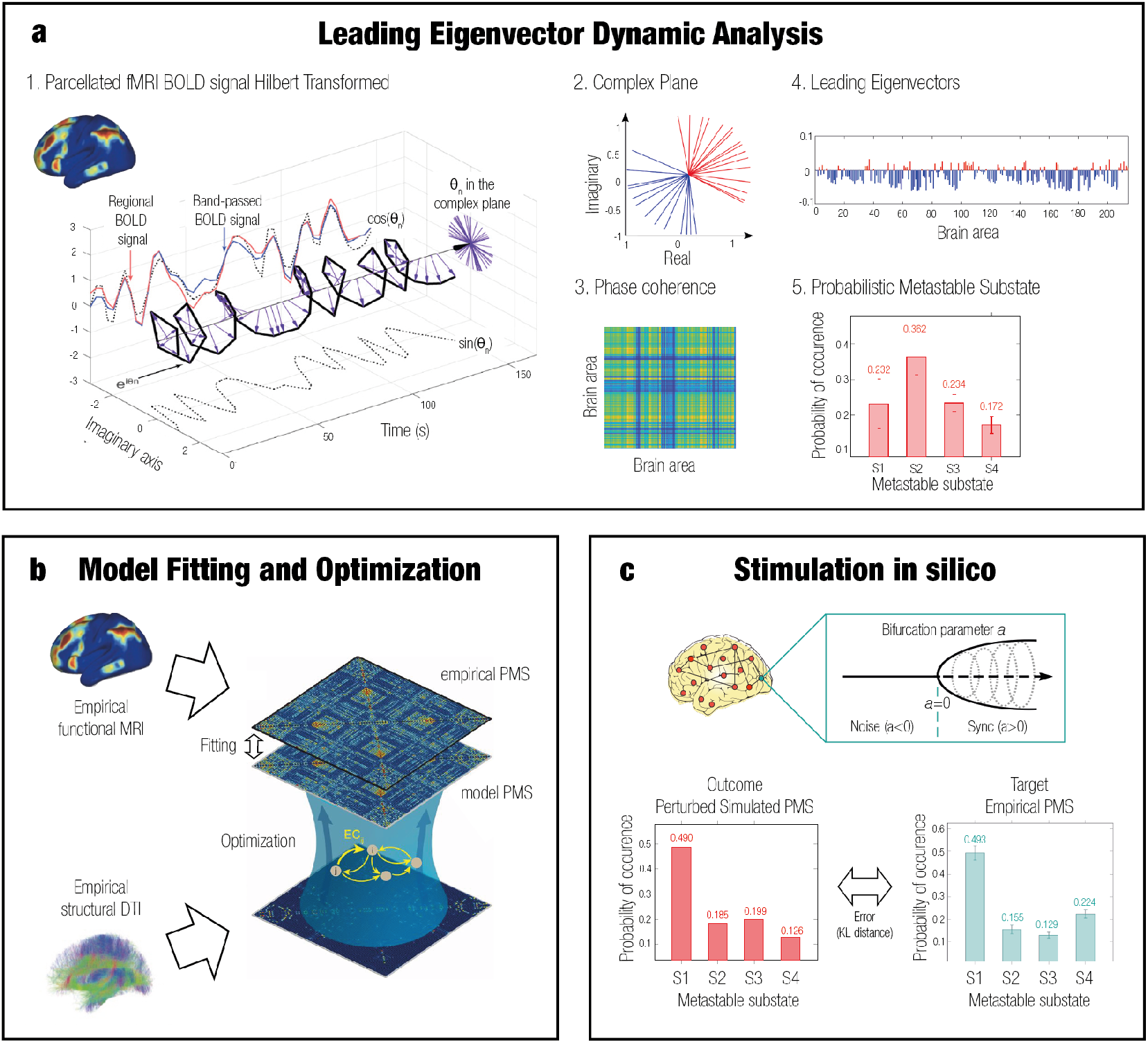
Overview of model-free and model-based frameworks. **a** Model-free framework: Leading Eigen-vector Dynamic Analysis (LEiDA). The BOLD time signal for each of the 214 brain areas was band-passed filtered and Hilbert transformed. The complex plane shows the positive and negative real and imaginary components at a specific timepoint *t*. The phase coherence matrix dFC(*t*) between brain areas for each time window was calculated. Then, the leading eigenvector *V*_1_(t) capturing the principal orientation of the BOLD phase for each of the matrices was calculated for each time *t* -positive values in red, negative values in blue. The leading eigenvectors for all time points of all participants were clustered using K-means (*k* =4), and the probability of occurrence of each of the cluster centers is shown in the Probabilistic Metastable Substate (PMS) Space. **b** Model-based framework: whole-brain model. A whole-brain model based on the frequency *w* of the empirical fMRI data and DTI was fitted to the empirical PMS space by calculating the value of the global coupling *G* that minimized the KL distance between the empirical and the simulated PMS. The model was optimized using the effective connectivity (EC) by adjusting each connection with a gradient descent approach until convergence. **c** Model-based framework: stimulation *in silico*. A transition was forced systematically from a source state to a target state by stimulating each brain area separately. The bifurcation parameter was shifted positively and negatively for synchronization and noise protocols, respectively. The optimal unilateral perturbation was obtained at the minimal KL distance between the stimulated modelled PMS and the target empirical PMS.

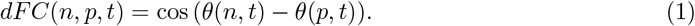

In this way, the interregional BOLD signal synchrony for all subjects was obtained at all time points. If nodes are temporarily aligned, the difference between their Hilbert transformed signal angle is 0° and the phase coherence is close to one [cos(0°)=1]. When a pair of nodes develop orthogonal BOLD signals, then the phase coherence is close to zero [cos(90°)=0]. The resulting dFC(*t*) of each subject at each timepoint was a 3D matrix of size of N*×*N*×*T, being N the number of brain areas (214) and T the total time points (295). A total of 69 3D matrices were calculated, corresponding to all of the groups together (controls, MCS and UWS).

In order to facilitate the future classification process, the dominant connectivity pattern was obtained by reducing the dimensionality of the matrices into their leading eigenvectors *V*_1_(t). This can be applied since FC matrices are undirected and symmetric across the diagonal (Deco et al., 2019). The leading eigenvectors (of dimension N*×*1) capture the dominant connectivity pattern at each time point *t* whilst explaining most of the variance, representing the contribution of each brain area to the whole structure and improving the signal-to-noise ratio (Cabral et al., 2017b). The dimensionality of the data was reduced from N*×*N to N*×*1, and the dominant functional connectivity pattern dFC(*t*) could be observed by calculating the outer product of *V*_1_(t) with its transpose (V1.V1.^*T*^) (Lohmann et al., 2010).

The following step consisted of identifying recurrent FC patterns representing the substates. The leading eigenvectors dFC(*t*) for each TR and all subjects from all states (20355 = 69 participants * 295 timepoints) were clustered with K-means clustering, varying *k* from 3 to 8. This algorithm is an unsupervised method consisting of assigning the data to the closest cluster centroid iteratively and re-calculating the *k* centroids in each iteration until convergence. The resulting cloud centroids *V*_*c*_(t) represent the dominant connectivity pattern in each cluster. The *k* discrete number of patterns of size N*×*1 correspond to the substates obtained from all subjects in all collapsed groups of subjects. These cluster centroids *V*_*c*_(t) represent the contribution of each brain area to the community structure and were rendered onto brain maps.

Upon computing the discrete number of FC patterns for each *k*, we calculated the resulting probability of occurrence in each group. This was computed as the ratio between the total number of epochs assigned to a specific cluster (i.e., for each subject in each group divided by the total amount of epochs in the given group). This gave rise to the Probabilistic Metastable Substate Space (PMS), which typifies each brain state from the probability of occurrence of being in each particular substate from the substate repertoire.

### Whole-Brain Computational Model

After characterizing the empirical PMS for the different profiles, a whole-brain Hopf computational model was obtained for each DoC state (**Figure 1b**). The dynamics from functional interactions between each brain area were emulated based on the anatomical SC. In other words, the emergence of activity can be explained in a mechanistic way by merging anatomical connectivity, which determines structure, and functional connectivity that represents activity dynamics, with the inclusion of effective connectivity (EC) (Deco et al., 2015). The working point of each model was fitted to the empirical data and optimized by determining the specific parameters of the model (Deco et al., 2019).

The normal form of supercritical Hopf bifurcation (Landau-Stuart oscillator) was used to simulate the BOLD activity for each of the 214 cortical and subcortical brain areas based in Shen parcellation. The Landau-Stuart oscillator has been used to study transitions from noisy to oscillatory regimes and, when coupled based on the brain’s architecture, to replicate complex interactions in brain dynamics (Deco et al., 2015).

An uncoupled node *n* can be represented in Cartesian coordinates with the following pair of coupled equations:

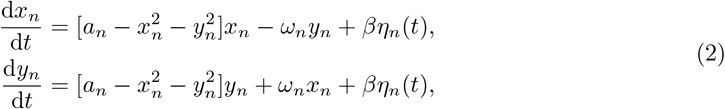

where *x*_*n*_ emulates the BOLD signal of the node and *η*_*n*_(*t*) is the additive Gaussian noise with standard deviation *β* = 0.01. This normal form describes the noisy and synchrony scenarios and has a supercritical bifurcation in *a*=0. For *a<0*, the node is stable in a fixed point and represented by noise from asynchronous firing of neurons. For *a>0*, metastable oscillations are obtained due to the synchronized firing of neurons at a frequency of *w/2π* (Deco et al., 2017b).*The transition from a noisy to a fully oscillatory scenario is called Hopf, and since it is the simplest way to model it mathematically, it is called normal form. Here, we chose a value of a_n_*=-0.02 for each brain node *n* following previous findings (Deco et al., 2017c), near the brink of Hopf bifurcation, in the critical border between synchrony and desynchrony. The frequency of the system *f*_*n*_ = *ω*_*n*_/2*π* was estimated from the empirical data as the averaged peak frequency of the filtered BOLD signal in the 0.04-to 0.07-Hz band for each brain node *n*=1,…, 214 (Deco et al., 2019).

The whole-brain dynamics were modelled by including an additive coupling term *C*_*np*_ which adjusts the input to node *n* from each of the rest of the nodes *p* based on the SC. This weighted matrix assumes different myelination densities across long-rage connectivities. A global coupling weight *G* was also added to represent the strength between all nodes, corresponding to the control parameter adjusted to fit the dynamical working region of the simulations to the empirical data. It scales all of the connections allowing maximal fitting between simulations and empirical data, assuming all axonal conductivity to be equal across the brain. The whole-brain dynamics at each node *n* was thus defined by the following set of coupled equations (Deco et al., 2017c):

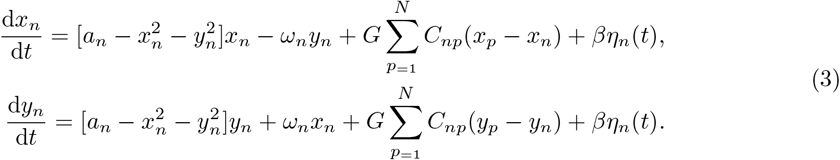

### Model Fitting: Comparing empirical and simulated probability metastable space states

For optimal spatiotemporal fit of whole-brain models to their empirical PMS space, the value of *G* was ranged from 0 to 0.5 in steps of 0.01, and the model was iterated 200 times. LEiDA was computed to the Hilbert-transformed simulated signal using the centroids already defined by the empirical substates in order to compute the simulated PMS space. Each model was fitted to the empirical data by deciding which value of *G* approximated it better (Deco et al., 2019). This corresponded to the lowest Kullback-Leibler (KL) distance between the empirical and simulated probabilities of each substate (Deco et al., 2019), given by:

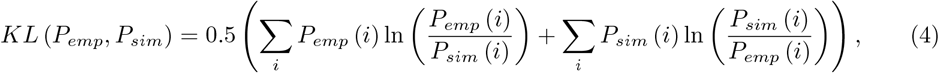

where *P*_*emp*_(*i*) and *P*_*sim*_(*i*) are the empirical and simulated probabilities respectively of metastable substate *i*.

### Model Optimization: Method for updating Effective Connectivity

After defining the value of *G* of each model, the models were optimized separately and the SC was updated in order to access potential missing connections. The initial value of *C* for each of the models was provided by a primer empirical DTI structural connectivity corresponding to the average of control subjects (Deco et al., 2015). Specifically, *C* was initially normalized to a maximum value of 0.2 in order to have the same range of values as in previous works (Deco et al., 2019, 2017c). The SC was then transformed to effective connectivity (EC) in an iterative manner by calculating the distance between the grand average phase coherence matrices of the model *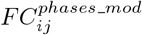* and the empirical matrices *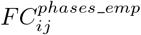*. Each structural connection between different nodes *i* and *j* was adjusted with a gradient descent approach given by:

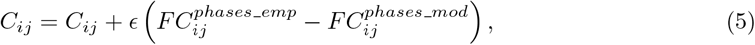

where *ϵ* = 0.01, and the grand average phase coherence matrices are defined:

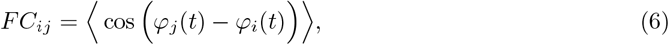

where *φ*(*t*) denotes the Hilbert transform BOLD signal phase of the nodes *j* and *i* at time *t*, and the brackets indicate the average across time. This was repeated until the difference between the empirical and simulated values was smaller than 0.001 (Deco et al., 2019)

### Unilateral Perturbation of the Whole-Brain Model

After obtaining the models, the transitions from the DoC states towards a control state were studied (**Figure 1c**). The models for DoCs were stimulated *in silico* by moving locally in a unilateral way the local bifurcation parameter *a* of each of the 214 brain areas. Different levels of intensity were applied area by area under the protocols of synchronization and noise. The protocols were represented by the sign of the local bifurcation parameter (positive and negative, respectively), and the stimulation intensities by the absolute value of each step (Deco et al., 2017c). In the synchronization protocol, the bifurcation parameter was shifted positively from 0 to 0.2 in steps of 0.01, whereas for the noise modality, it was shifted from 0 to -0.2 in steps of -0.02. Each simulation was repeated 3 times the results were averaged to minimize random effects from the Gaussian noise of the model (Deco et al., 2019). The fitting to the target states was measured by calculating the KL distance (described in the previous section) between the probabilities of each substate of the simulated DoC models separately, which are the source, and the empirical control PMS, which is the target. The areas more prone to promote a desired transition after simulation were detected from the ones presenting the lowest KL distance.

### Statistical Analysis

Statistical analysis were performed using MATLAB R2022a software from MathWorks (Natick, MA, USA). Permutation-based Wilcoxon tests with 1000 iterations were used to test the results of the LEiDA method, specifically the probability of occurrence of the whole range of explored clustering conditions (*k* from 3 to 8). The Wilcoxon test was used to compare each permutation with a significance threshold of 0.05. We applied the False Discovery Rate (FDR) method (Hochberg and Benjamini, 1990) to correct for multiple comparisons when testing the differences between groups (controls, MCS, and UWS) and the number of cluster centers (i.e., substates). All p-values shown correspond to the differences that remain significant after FDR correction.

## Results

### LEiDA

We selected the minimum number of clusters (*k*) that statistically differed between the three groups. The configuration that best described the empirical data across all participants and distinguished between groups was detected at *k* =4. The probability of occurrence for the PMS of each group is visualized in **Figure 2a** and the cluster centroid eigenvectors are rendered onto brain maps in **Figure 2b**. The leading eigenvectors had positive and negative signs partitioning the network into communities as red and blue colors, respectively. The strength of the color describes the strength with which each area belonged to the placed community (Cabral et al., 2017b).

**Figure 2:**
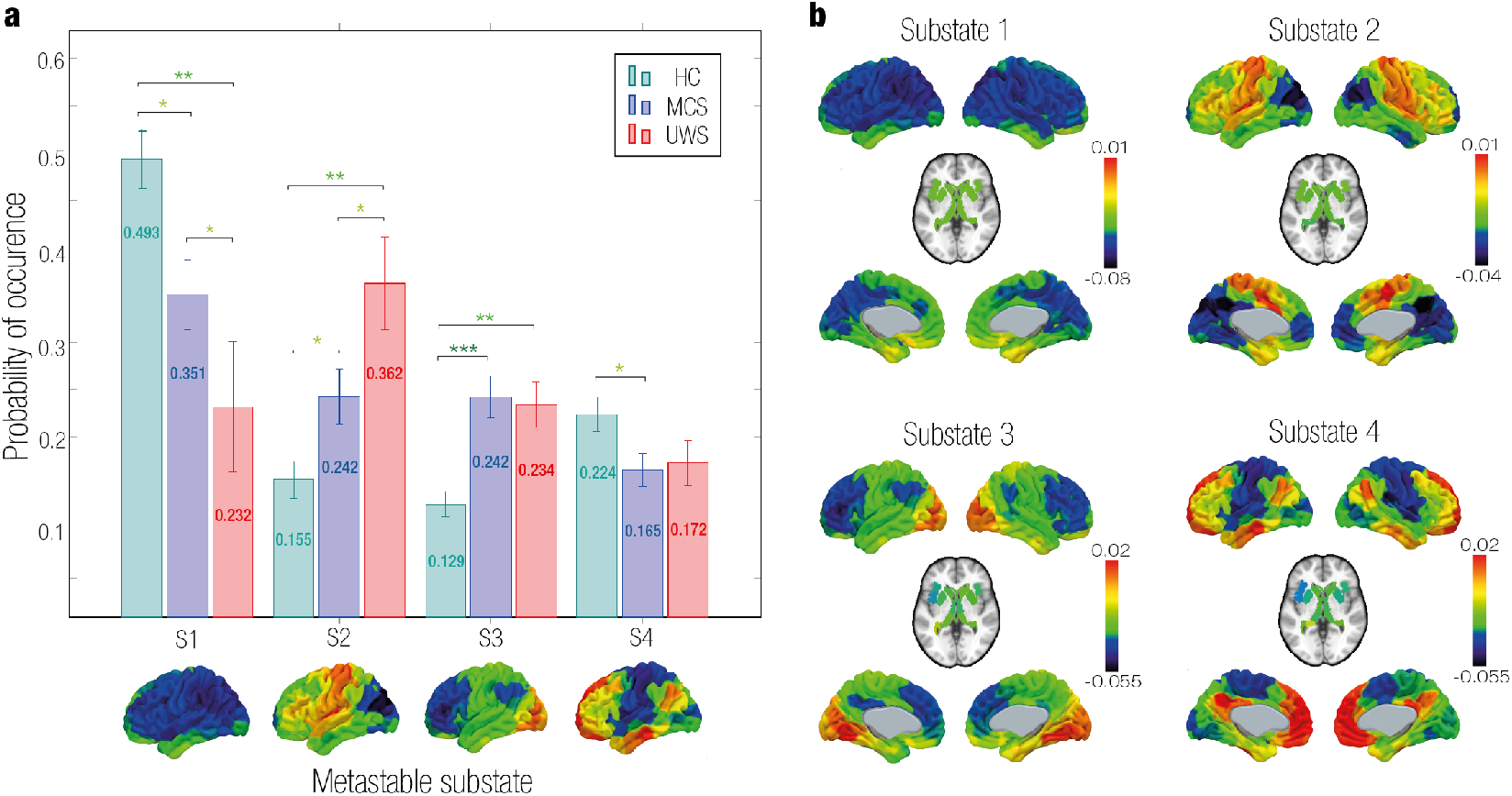
Model-free results: Empirical Probabilistic Metastable Substate (PMS) Space. **a** Probability of Occurrence. The mean probability of occurrence for each group in each substate was calculated with a 95% confidence interval. The substates 1 and 4 had a higher probability of occurrence for the control group compared to DoC. The substates 2 and 4 had a lower probability of occurrence for the control group compared to DoC. Statistically significant differences are represented with asterisks (* p *<* 0.05, ** p *<* 0.01 and *** p *<* 0.001). **b** Rendered brains represent the leading eigenvectors of each substate plotted onto the cortex. Substate 1 was characterized by all elements of the eigenvector with the same sign. Substate 2 had a functional community formed by areas in the motor network. Substate 3 presented a local coordination in the occipital lobe (visual network). Substate 4 showed coordination in brain areas from the medial-frontal network, fronto-parietal network, DMN and subcortical areas.

The first substate presented the same sign for all eigenvector elements. The probability of occurrence was higher in controls [0.493 ± 0.030 (mean ± standard error)] compared to MCS [0.351 ± 0.037, *P* =0.012] and UWS [0.232 ± 0.069, *P* =0.003]. Furthermore, the probability was lower in UWS than in MCS [*P* =0.035]. The rest of the substates (i.e., substates 2, 3, and 4) were characterized by subsets of brain areas that disengaged from the whole-brain network aligning with each other. In substate 2, central areas (motor network) represented a pattern of activation. In this substate controls had the lowest probability of occurrence [0.155 ± 0.020] compared to MCS [0.242 ± 0.029, *P* =0.022] and UWS [0.362 ± 0.050, *P* =0.001]. Moreover, the probability was higher in UWS than in MCS [*P* =0.033]. Substate 3 exhibited a functional network led by the occipital lobe (visual network). In controls, the probability of substate 3 was lowest [0.129 ± 0.013] compared to MCS [0.242 ± 0.022, *P<*0.001] and UWS [0.234 ± 0.024, *P* =0.001]. This substate did not discriminate significantly between DoC groups. Substate 4 had a coordination between areas of the medial-frontal network, fronto-parietal network, DMN (i.e., precuneus) and subcortical areas (i.e., thalamus). This metastable substate only discriminated between controls [0.224 ± 0.018] and MCS [0.165 ± 0.018, *P* =0.018].

### Fit whole-brain computational model to the brain states of DoC groups

For the MCS and UWS groups, we fitted the PMS to a causal mechanistic whole-brain model. We optimized and adjusted the models in order to select the parameters that displayed the most approximate regime to empirical PMS (see Materials and Methods). The best fit between the empirical and simulated PMS was found at *G* =0.08 and *G* =0.05 for MCS and UWS models, respectively (**Figure 3**).

**Figure 3:**
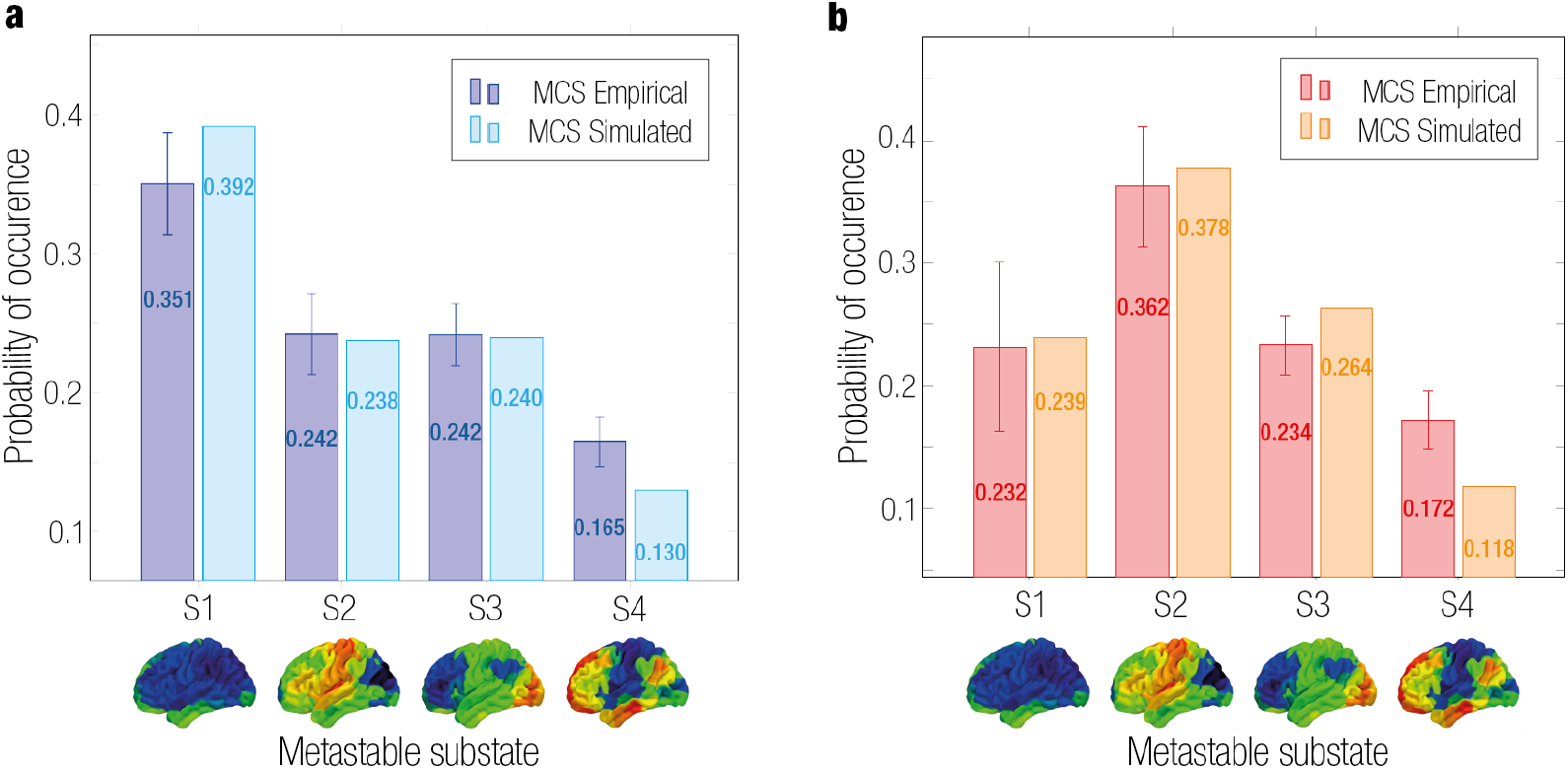
Model-based results: Whole-brain model fitting and optimization. Comparison between empirical and simulated PMS of each group. Optimal fit was given by the minimal KL distance value corresponding to a global coupling weight of **a** *G*=0.08 for MCS and **b** *G*=0.05 for UWS.

### *In silico* stimulations to force transitions from DoC to a control target state

Following model fitting and optimization, we systematically perturbed the PMS model of each DoC group and compared it with the empirical PMS of the control group. Each brain node was shifted by increasing the absolute value of the bifurcation parameter *a*, representing the intensity of stimulation. A synchronization protocol was addressed with positive values, and a noise protocol with negative values. Optimal perturbation was the one that resulted in the smallest KL distance between the PMS after perturbing each node individually, and the empirical PMS of the target (control group).

The results of the *in silico* stimulation with different protocols and intensities for MCS and UWS are shown in **Figures 4a** and **4b**, respectively. The color scale represents the KL distance between the perturbed PMS and the target PMS after stimulating each individual brain area separately. The best fit is indicated by a lower KL distance, note that the color scales are different for each DoC condition, adjusted accordingly for better resolution. For the synchronization protocol, a successful transition was forced from the source states of MCS and UWS to the control state. We can observe that for MCS most regions promoted a transition with lower stimulation intensity compared to UWS. In contrast, in the noise protocol, the KL distance did not decrease for both MCS and UWS (i.e., colors are red and yellow rather than green and blue). This means that as a result of applying a noise protocol, the transition from DoC to a control target state was not possible, evidenced by poorer fit.

**Figure 4:**
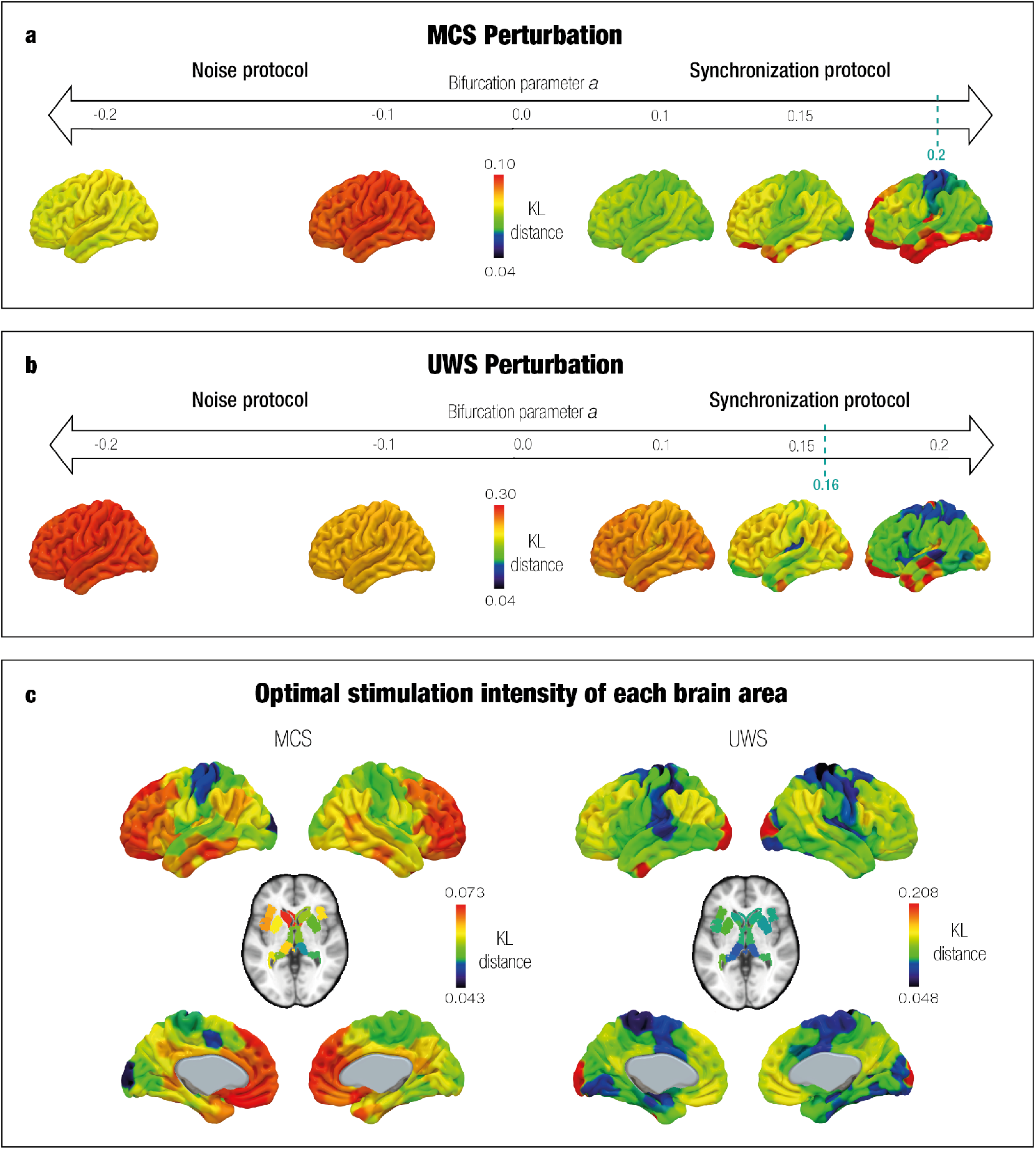
Model-based results: *In silico* probing to force transition from DoC to control target state. We used synchronization and noise stimulation protocols to shift the local bifurcation parameter. The strength of the unilateral perturbation corresponds to the absolute value of the bifurcation parameter and the sign to the modality (synchronous with positive values, noise with negative values). The x-axis shows the stimulation intensity (from softer to stronger), and the color scale represents the KL distance. The best effectiveness was found where KL distance was minimal. For both DoC (**a** and **b**), the synchronization protocol forced a transition to the control state. This can be observed with the lower KL distance when increasing values of the local bifurcation parameter in a positive manner. The left sides of the x-axis show that the noise protocol presented poor effectiveness given that KL distances were longer than in the synchronization protocol. **c** contains the KL distance rendered onto brain maps with the optimal stimulation for each brain area in the synchronous protocols. The color scale represents the KL distance given by the best stimulation, with the lowest values corresponding to the motor and some subcortical areas (the best targets).

In the synchronization protocol, a transition was likely to occur in many areas if they were sufficiently stimulated. **Figure 4c** illustrates the rendering of the KL distance between the perturbed PMS and the target PMS after stimulating each individual brain area separately, at their particular optimal stimulation intensity. Areas in the motor network were the most sensitive ones, including subcortical areas (i.e., thalamus), provoking transitions in both cases (MCS and UWS). Specifically, the best fit to the control PMS space was obtained when stimulating the left post-central gyrus with an intensity of 0.2 for MCS and the right postcentral gyrus with an intensity of 0.16 for UWS. As a result, the perturbed and target probabilities were very similar in all four metastable substates of the PMS **(Figure 5)**.

**Figure 5:**
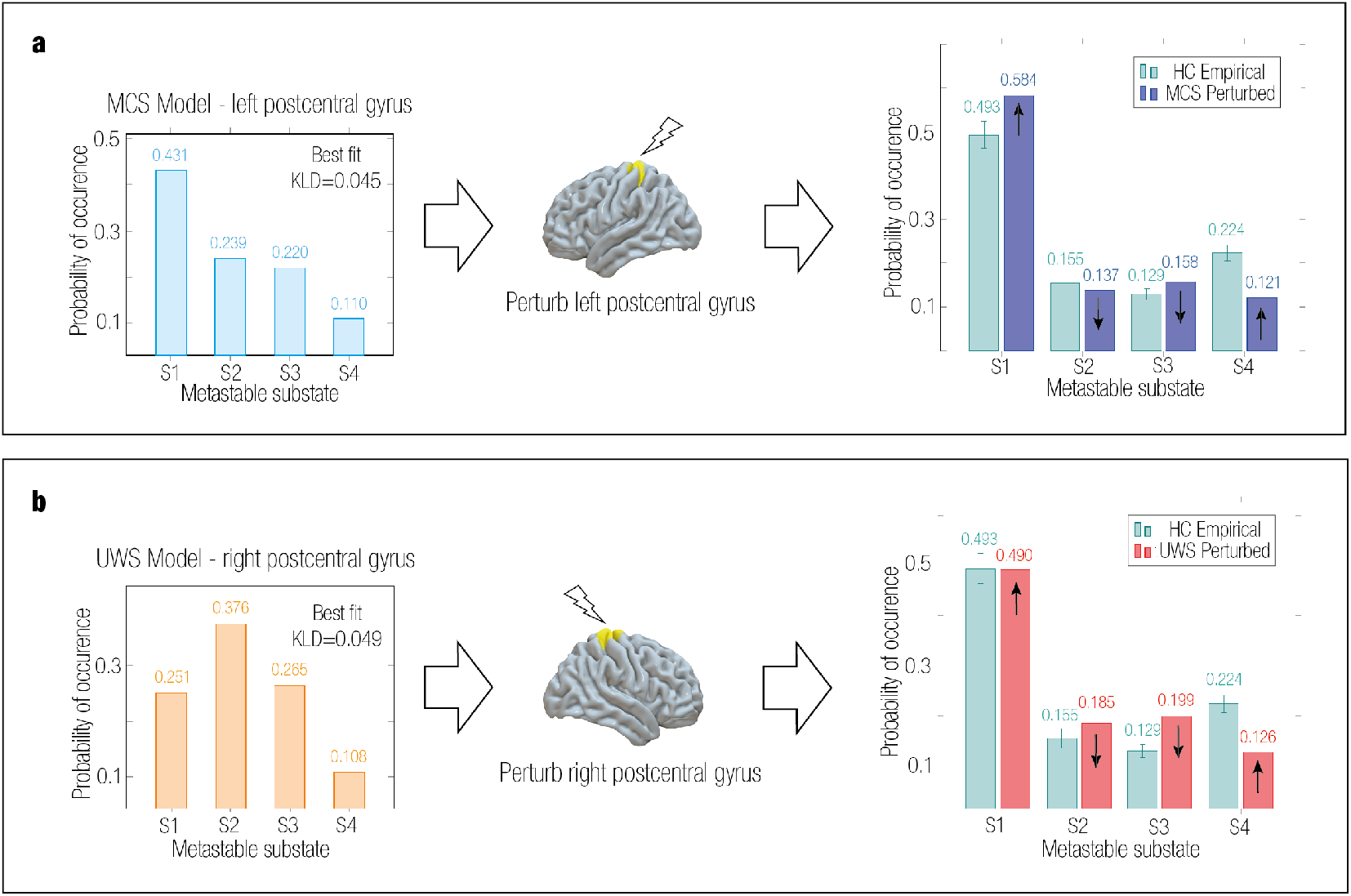
Comparison between perturbed PMS of MCS and UWS groups to target control PMS. We show the simulated and perturbed PMS for DoC groups and the empirical target control PMS. For both groups, the synchronization protocol increased the probability of the first and last substates and decreased the probability of the other substates, consistent with the empirical PMS of the control group. **a** Simulated MCS had a best approximation to the PMS of controls by perturbing unilaterally the postcentral gyrus (left) with an intensity of 0.2. **b** Simulated UWS had the best approximation to the PMS of controls by perturbing unilaterally the postcentral gyrus (right) with an intensity of 0.16.

## Discussion

We successfully applied model-free and model-based approaches to find causal evidence for the brain dynamics in DoC and transitions to a control state, following the methodology of Deco et al. (2019). Firstly, we significantly distinguished between brain states by characterizing the PMS of DoC and controls using LEiDA. For each group, we identified metastable substates (i.e., patterns) with an associated probability of occurrences and alternation profiles (Cabral et al., 2017b). We then fitted a Hopf model to each empirical PMS for each DoC state. In this way, we were able to force a transition from the PMS DoC models (MCS and UWS separately) to the target control state using exhaustive off-line *in silico* unilateral perturbations. Finally, by varying stimulation intensities, we revealed how changes in local brain areas using a synchronous modality can reshape whole-brain dynamics in DoC. In this way, we could determine the mechanistic global effects of all possible local perturbations and the most sensitive areas in terms of their perturbability.

In the model-free approach, using LEiDA, we identified substates with network-specific changes whose probabilities varied in each brain state (**Figure 2**). In particular, we found controls were more able to access substates 1 and 4. Substate 1, in which all BOLD signals followed the leading eigenvector, has been shown to exist in previous LEiDA studies (Lord et al., 2019; Figueroa et al., 2019; Cabral et al., 2017a). This substate has been associated with a global state (Zhang and Northoff, 2022), synchronized stability (Farinha et al., 2022), or noise artifacts (Olsen et al., 2022). Furthermore, we found substate 4 had a coordination of areas overlapping the medial-frontal network, fronto-parietal network, DMN (i.e., precuneus) and subcortical areas (i.e., thalamus). The DMN is important for internal self-related and external perceptual awareness, cognition, mind-wandering, and autobiographical memory, and some studies have shown this network disrupted in patients with DoC (Panda et al., 2022; Edlow et al., 2020; Bodien et al., 2017; Qin et al., 2015; Demertzi et al., 2015; Fernández-Espejo et al., 2012; Vanhaudenhuyse et al., 2011, 2009; Demertzi et al., 2013). On the other hand, our results show DoC patients were more likely to be in substates 2 and 3, which exhibited a functional network led mainly by areas from the motor and visual networks, respectively. Notably, Demertzi et al. (2015) found a correlation between rs-fMRI connectivity of the aforementioned networks (DMN, fronto-parietal, sensorimotor, and visual networks) and CRS-R assessment results, indicating these networks are critical to brain function in DoC. This is also supported by Cao et al. (2019), which reported changes in brain activity in the DMN, somatomotor, and visual networks, and by Crone et al. (2014), which measured altered network properties in the fronto-parietal cortex, both studies in DoC patients.

In the model-based approach, we modelled brain activity as a system of non-linear Stuart-Landau oscillators (also known as Hopf bifurcation) to link the underlying anatomy with local dynamics (Deco et al., 2017c). Hopf models have allowed simulating several brain states in health and disease with high fitting accuracy (Escrichs et al., 2021b, 2022; Sanz Perl et al., 2023; Soler-Toscano et al., 2022; López-González et al., 2021; Deco et al., 2019; Jobst et al., 2017; Deco and Kringelbach, 2014). These models have been able to capture both local and global brain dynamics (Deco et al., 2017c,b), while having lower computational costs (Deco et al., 2017a; Deco and Jirsa, 2012) and risks of overfitting (Deco and Kringelbach, 2014) than more detailed models such as spiking neurons (Deco and Jirsa, 2012; Cabral et al., 2014). Here, during fitting and optimization, we observed that the MCS group had a higher value of global coupling weight *G* than the UWS (**Figure 3**). This parameter represents the relationship between local and global brain dynamics and the effects of structural connectivity on brain dynamics. The greater the value of *G*, the less restricted the brain network interaction is to areas with high structural connections. In line with previous studies, we found that MCS showed more propagation of brain activity and connectivity between distant brain areas than UWS (Escrichs et al., 2021b; López-González et al., 2021).

By combining the model-based approach with *in silico* stimulations, we explored brain transitions between different states. This strategy allowed us to find the optimal areas to stimulate and re-balance the underlying brain dynamics in patients with DoC towards more healthy states (Escrichs et al., 2021b; Sanz Perl et al., 2021). Thus, *in silico* stimulation provided us a way to test exhaustive trials without the ethical constraints of real-world experiments (Deco et al., 2017c; Clausen, 2010). We shifted the brain dynamics’ landscape rather than the working point per se. This ensures propagation and facilitates plasticity, targeting a system reorganization with long-term effects (Deco et al., 2019, 2017a). We evidenced transition from DoC states to control using the synchronization and not the noise protocol, in line with Deco et al. (2019), since the KL distance between the perturbed PMS and the control target PMS decreased in the synchronous modality (**Figure 4**). Bifurcation parameters below the bifurcation edge were, therefore, indicative of DoC states and could not force systems to a control target. Both MCS and UWS progressively came closer to the target state with increasing positive intensities for the desired transitions (i.e., the synchronization protocol). In this regard, our results are consistent with the notion that synchronous oscillations have a role in neuronal communication and long-range functional connectivity between brain areas (Cabral et al., 2022; Fries, 2005). A further finding was that overall, MCS exhibited higher sensitivity to external perturbations than UWS. Metastable substates with the highest probability of PMS spaces in both DoC groups shifted from substates with subsets of brain areas aligned within each other in the motor network (substate 2) and visual network (substate 3) to a substate dominated by regions from the medial-frontal network, fronto-parietal network, DMN and subcortical areas (substate 4), and to a substate with global brain activity (substate 1) (**Figure 5**). In terms of brain areas promoting a transition, most were found in the motor network, relevant to DoC (Panda et al., 2023; López-González et al., 2021; Demertzi et al., 2015; Piccione et al., 2011). Particularly, the most sensitive area was the postcentral gyrus, which has been associated with impaired somatosensory functions (Cao et al., 2019) and found to distinguish DoC patients by its weighted global connectivity (Kotchoubey et al., 2013). Lastly, a specific subcortical area prone to transition and important in DoC studies was the thalamus (Panda et al., 2023, 2022; Sanz Perl et al., 2021; Lutkenhoff et al., 2015; Monti et al., 2015; Schiff et al., 2007), given its key role in information processing and as a sensory relay station (Alnagger et al., 2023; Zheng et al., 2021).

The classification of patients with DoC is an existing debate in neuroscience. Identifying MCS and UWS can depend on the CRS-R metric’s effectiveness, inter-rater variability, and consistency of caregivers’ reports (Opara et al., 2014). It is challenging to distinguish between MCS and UWS since some patients who are classified as UWS may remain aware even though they do not demonstrate behavioral signs. They may be classified incorrectly as being awake and unaware when they are actually conscious (Owen, 2020; Bodien et al., 2015; Tagliazzucchi and Laufs, 2014; Fingelkurts et al., 2014). The circular nature of brain state definition and assessment could have compromised the efficacy and validity of our model definitions since they are subjected to the correct classification and typification of the empirical primary data source (Arsiwalla and Verschure, 2018). It would be helpful to investigate the generalizability of our results with a broader range of DoC patients (Vohryzek et al., 2022a; López-González et al., 2021).

Overall, we were able to characterize and differentiate brain dynamics of DoC and healthy controls. We used a robust quantitative definition of brain states based on spontaneous spatiotemporal fluctuations (Deco et al., 2015; Constable, 2006). Furthermore, we provide a causal mechanistic explanation for the differences between brain states in DoC. Crucially, our perturbation approach could be used as a specific model biomarker relating local activity with global brain dynamics. In light of the exciting results, future applications could benefit from developing personalized protocols by constructing individualized patient brain models (Vohryzek et al., 2022a,b; Kringelbach and Deco, 2020; Luppi et al., 2019; Muldoon et al., 2016; Constable, 2006). In addition, causal whole-brain modelling can help understand other brain states (e.g., meditation, anesthesia) (Seth and Bayne, 2022) and elucidate propagation properties (Rossini et al., 2015), network level impact (Kringelbach and Deco, 2020; Muldoon et al., 2016) and sensitive areas (Ipiña et al., 2020; Kringelbach et al., 2011). Overall, our results may eventually contribute to the field of external perturbation as a principled way of re-balancing the dynamics of post-coma patients towards more healthy regimes.

## Data availability statement

Due to the restrictions imposed by the approved ethics protocols, neuroimaging datasets cannot be shared publicly since they contain clinical information from patients. However, the data can be requested to the authors.

## Funding

P.D. was supported by the FI-SDUR Grant (no. 2022 FISDU 00229) funded by the Catalan Agency for Management of University and Research Grants (AGAUR). A.E. and G.D. were supported by the project eBRAIN-Health -Actionable Multilevel Health Data (id 101058516), funded by the EU Horizon Europe. G.D. was also supported by the AGAUR research support grant (ref. 2021 SGR 00917) funded by the Department of Research and Universities of the Generalitat of Catalunya and by the project NEurological MEchanismS of Injury, and the project Sleep-like cellular dynamics (NEMESIS) (ref. 101071900) funded by the EU ERC Synergy Horizon Europe. Y.S.P. was supported by the European Union’s Horizon 2020 research and innovation program under the Marie Sklodowska-Curie grant 896354. S.L., and J.A. were supported by the HBP SGA3 Human Brain Project Specific Grant Agreement 3 (grant agreement no. 945539), funded by the EU H2020 FET Flagship. The study was supported by the University and University Hospital of Liège, the Belgian National Funds for Scientific Research (F.R.S-FNRS), the MIS FNRS project (F.4521.23), the BIAL Foundation, AstraZeneca Foundation, the Generet funds and the King Baudouin Foundation, the James McDonnell Foundation, and Mind Science Foundation. O.G. is a research associate and S.L. is a research director at the F.R.S-FNRS.

## Competing interests

The authors declare no conflict of interest.

## Supporting information

**S1 Table. Top 20 most sensitive regions for perturbing MCS model.** The first column corresponds to the KL distance between the PMS of the perturbed model, and the PMS of the target control state, after stimulating a given brain area. The second column shows the brain area in the Shen parcellation (Shen et al., 2013). The third column indicates the overlap between the brain area and the AAL structural parcellation (Tzourio-Mazoyer et al., 2002).

**S2 Table. Top 20 most sensitive regions for perturbing UWS model.** The first column corresponds to the KL distance between the PMS of the perturbed model, and the PMS of the target control state, after stimulating a given brain area. The second column shows the brain area in the Shen parcellation (Shen et al., 2013). The third column indicates the overlap between the brain area and the AAL structural parcellation (Tzourio-Mazoyer et al., 2002).

## Notes

### Competing Interest Statement

The authors have declared no competing interest.

